# Reaction-Conditioned Enzyme Discovery with Multimodal Deep Learning

**DOI:** 10.64898/2026.03.09.710689

**Authors:** Ziyi Zhou, Yutong Hu, Yuanzhen Zhang, Xinnan Fu, Runye Huang, Bozitao Zhong, Xiaoran Cheng, Jin Huang, Qin Xu, Shuangjun Lin, Linquan Bai, Liang Hong, Pan Tan

**Affiliations:** Shanghai National Center for Applied Mathematics (SJTU Center) & Institute of Natural Sciences, Shanghai Jiao Tong University, Shanghai, China; School of Computer Science, Shanghai Jiao Tong University, Shanghai, China; Shanghai Artificial Intelligence Laboratory, Shanghai, China; State Key Laboratory of Microbial Metabolism, and School of Life Sciences & Biotechnology, Shanghai Jiao Tong University, Shanghai, China; Core Facility and Service Center (CFSC) for School of Life Sciences & Biotechnology, Shanghai Jiao Tong University, Shanghai, China; Zhangjiang Institute for Advanced Study, Shanghai Jiao Tong University, Shanghai, China

## Abstract

The precise mapping between chemical transformations and enzymatic catalysts underpins the complexity of metabolic networks. Conventional discovery methods, tethered to sequence homology or structural alignment, are inherently blind to new reactions. Here we present VenusRXN, a multimodal deep learning framework that shatters this limitation by enabling reaction-conditioned enzyme discovery. By seamlessly unifying a pre-trained reaction encoder with a protein language model, VenusRXN achieves a fine-grained, high-dimensional alignment of chemical and biological representations. On benchmarks to discover enzymes which catalyze reactions not seen in the training dataset, it surpasses state-of-the-art baselines with a top-20 retrieval hit rate of 76.5%. Most critically, we demonstrate VenusRXN’s capabilities in a zero-shot discovery. As verified by the wet-lab experiments, it successfully identified enzymes to catalyze the chemical reactions never reported, including the one to catalyze the synthesis route for a type 2 diabetes drug intermediate using a non-natural substrate. With surprising precision, the model pinpointed active candidates within the top 10 sequences directly from a global search space of over 300 million proteins, which can hardly be achieved by structure-based enzyme discovery algorithm. Thus, VenusRXN unlocks the capacity to interrogate the vast, unannotated “dark matter” of the protein universe with affordable computational cost for everyone. This work signals a definitive paradigm shift, establishing the chemical reaction itself, rather than homology, as the primary functional descriptor for the *de novo* discovery of biocatalysts.

## Introduction

The relationship between diverse chemical reactions and highly specific enzymes shapes the complex architecture of metabolic networks. The diversity of enzymatic reactions introduces alternative and bypass routes, rendering the metabolic network branched and modular, while enzyme reaction specificity constrains flux via substrate and stereochemical selectivity. Together, these features balance the robustness and precision of metabolism. Since less than 1% of the billions of sequenced proteins are functionally characterized [1], establishing accurate reaction-enzyme mapping, i.e., accurately identifying enzymes with specific catalytic functions, is a foundational prerequisite for function-driven enzyme discovery and an urgent need for many applications [2]. This is particularly critical in natural product biosynthesis, where such mapping is indispensable for the discovery [3], annotation [4], and functional reconstruction [5] of biosynthetic gene clusters.

However, conventional homology-based enzyme function annotation methods struggle to predict reaction-enzyme mapping directly and accurately. They typically infer enzyme function by searching for sequence or structural homologs of the query protein in annotated databases using tools like BLASTp [6] and Foldseek [7], then transferring the annotation of the best match. Despite being widely used, the reliability of such approaches is limited, as similarity in protein sequence or structure does not always guarantee functional consistency. Recently, there has been a surge in deep learning-based enzyme function prediction methods. They leverage protein language models (PLMs) [8-10] to predict functional descriptors such as enzyme commission (EC) number [11-13] and gene ontology (GO) terms [14,15]. Yet, they are criticized by their high false positive rates and limited interpretability [16]. Most importantly, because all the above methods rely on existing annotations, they remain incapable of handling reactions that are new-to-nature or beyond the scope of current annotation systems.

Therefore, reaction-conditioned enzyme discovery represents a more promising paradigm, in which target enzymes can be found directly from their catalyzed reactions, which in principle can discover novel enzymes to catalyze reactions not reported by using biological catalysis. As a step towards this goal, a line of research pursues to model the substrate-enzyme mapping [17-19], but focusing on the substrate alone captures only part of the reaction and leaves products unspecified. Although several models have been proposed to directly learn reaction-enzyme mapping [20-23], their performances remain limited and they still face tough challenges: (1) the absence of robust reaction encoding models for high-quality reaction representations; (2) the need for effective training strategies to learn complicated cross-modal interactions between reactions and enzymes.

To address the above challenges, we introduce VenusRXN, a reaction-conditioned enzyme discovery framework based on multimodal deep learning. In VenusRXN, we develop a molecular graph-based reaction encoder to extract robust reaction representations. By leveraging self-supervised pretraining and condensed graph of reaction (CGR) [24], the reaction encoder effectively models the structural transformations from reactants to products. To learn reaction-enzyme mapping, VenusRXN integrates the reaction encoder with a protein language model (PLM), and jointly trains them on a large reaction-enzyme dataset through multimodal learning. The joint training objectives involve contrastive learning [25], soft-label alignment [26], and classification of reaction-enzyme pairs, enabling fine-grained matching and fusion of reaction and enzyme representations. Once trained, VenusRXN supports three applications: retrieval of enzymes that can catalyze the given query reaction, retrieval of enzymes that have similar catalytic function to the given template enzyme, and fine-tuning using catalytic performance labels for task-specific enzyme recommendation.

Extensive benchmarking reveals that VenusRXN surpasses state-of-the-art methods, establishing a new frontier in zero-shot generalization for reactions entirely absent from the training data. On our curated benchmark, the model achieves a top-20 retrieval hit rate of 76.5% for reactions unseen during training. Furthermore, VenusRXN accurately reconstructed four recently elucidated natural product biosynthetic pathways (postdating the training dataset) through genome-wide screening, ranking all target enzymes within the top 50 predictions. This capability stands in stark contrast to traditional elucidation strategies that rely on laborious gene knockouts and complex heterologous co-expression assays, offering a streamlined computational solution that dramatically reduces the experimental burden.

Crucially, unlike existing methods that depend on protein structures [21,22] or pre-defined pocket information [19], VenusRXN shatters these scalability barriers by learning a direct, sequence-exclusive mapping between chemical logic and protein function. This architectural distinction is paramount: as the explosive trajectory of genomic sequencing leaves structural data orders of magnitude behind, reliance on structure restricts discovery to a shrinking fraction of the biological landscape. In contrast, by constructing a high-dimensional vector index of the proteome, VenusRXN enables the rapid, end-to-end interrogation of the massive, unannotated “dark matter” within sequence databases. Leveraging VenusRXN, we constructed a vector database capable of retrieving enzymes from billion-scale protein sequences within minutes. This allows for the design of enzymes with state-of-the-art accuracy across the vast protein universe at a low computational cost affordable for everyone.

To rigorously validate this zero-shot capability in a hundreds of million-sequence search space, we challenged VenusRXN to perform template-free enzyme mining directly from the >300 million sequences in the NCBI non-redundant (NR) database [27]. Specifically, we define zero-shot as the capability to handle reactions and substrates entirely absent from the training data, and template-free as the ability to identify catalysts without reliance on known protein templates. We targeted two chemically complex transformations pivotal to type 2 diabetes therapeutics. These include a previously unreported reaction involving a non-natural substrate for glip class drug. With surgical precision, VenusRXN identified high-activity biocatalysts within the top 10 ranked candidates out of hundreds of millions. This “finding a needle in a haystack” success establishes VenusRXN as an advanced engine for zero-shot biocatalyst discovery, unlocking the boundless potential of new-to-nature biocatalyst.

## Results

### Framework of VenusRXN

A critical foundation for reaction-conditioned enzyme discovery is to obtain high-quality reaction representations capturing important atomic motifs and chemical rules in reactions. Therefore, we develop a reaction encoder that takes molecular graphs as input and encodes reactions from the atomic scale. Its core components include a pretrained Graph Transformer (Graphormer) model named Mol-Graphormer. Given a reaction, Mol-Graphormer encodes its reactant and product graphs separately to provide basic molecular features. Mol-Graphormer extends the sequence self-attention in standard Transformer blocks to graph self-attention, introducing spatial encoding and edge encoding to capture structural relation between atoms (Fig. 1a and Methods). We pretrain Mol-Graphormer on chemical reactions through self-supervised learning, where three reaction-aware pretraining objectives are designed (Fig. 1b and Methods): masked atom property prediction (MAP), reaction center prediction (RCP), and graph contrastive learning (GCL). These objectives help the model to learn contextualized representations of graph nodes, localize key atoms and functional groups involved in the reactions, and capture general reaction rules. To better reflect the structural changes from reactant(s) to product(s) in a reaction, the reactant and product graphs encoded by Mol-Graphormer are superposed based on atom mapping to construct a CGR [24]. Then, another Graphormer named CGR-Graphormer is applied to encode the CGR, and provides the final, high-level reaction representation (Fig. 1c and Methods). Its model architecture is similar to that of Mol-Graphormer but it is much shallower.

**Fig. 1.**
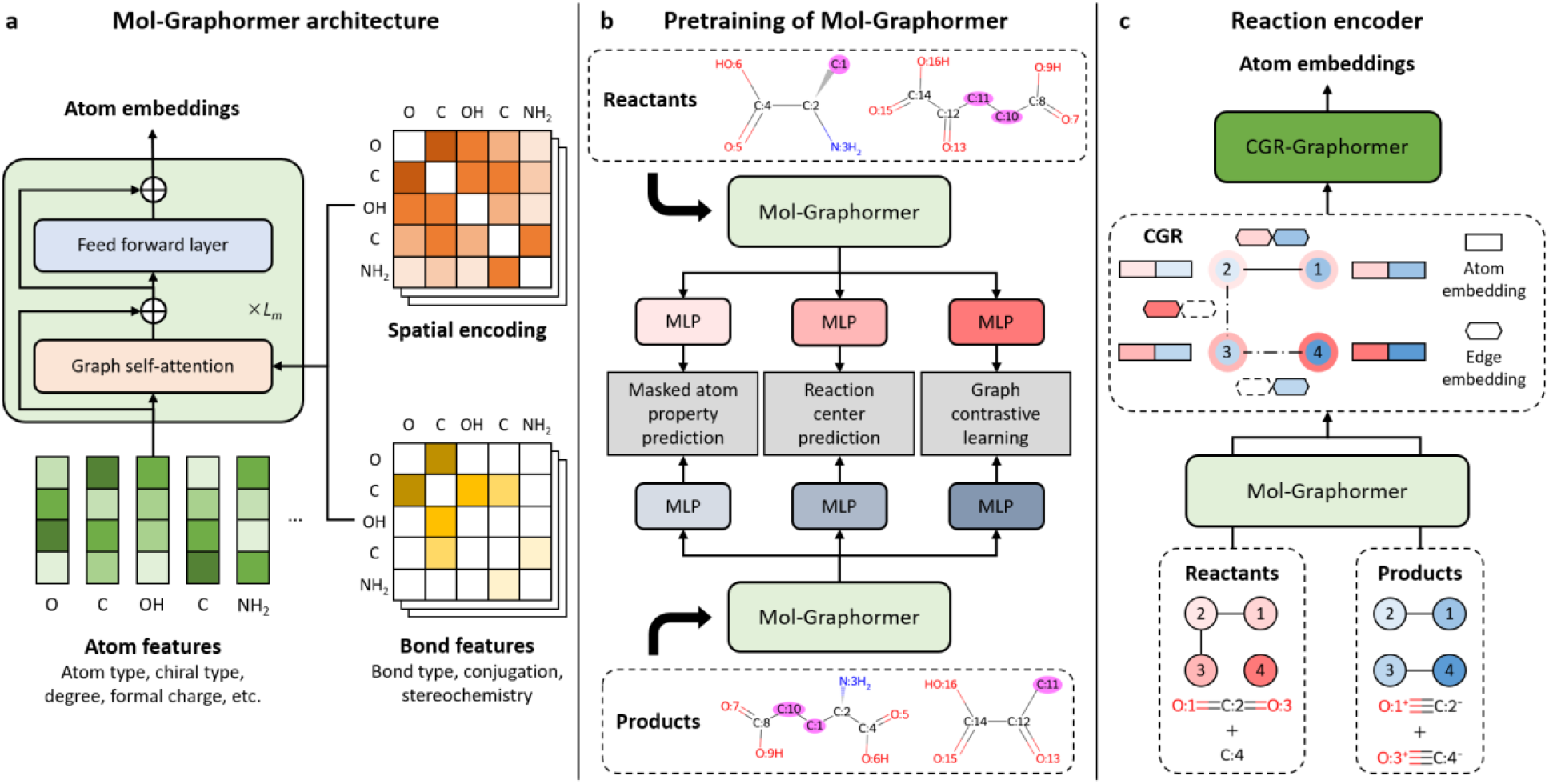
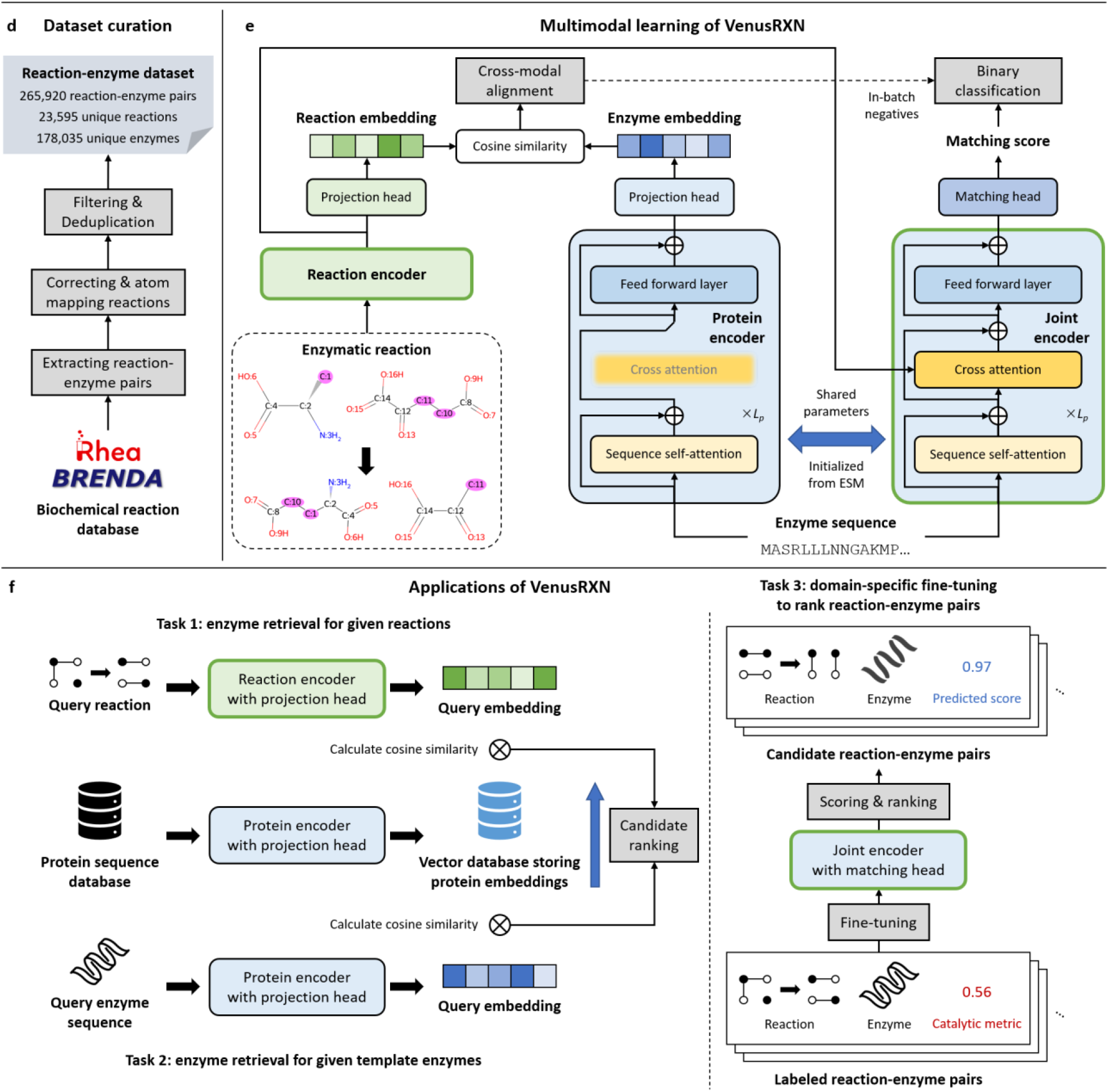
Framework of VenusRXN. **a** Mol-Graphormer aims to extract fundamental molecular graph features of reactants or products. It employs graph self-attention to capture structural relation between atoms. **b** Mol-Graphormer is pretrained on chemical reactions through three reaction-aware objectives: masked atom property prediction (MAP), reaction center prediction (RCP), and graph contrastive learning (GCL). **c** The reactant and product graphs encoded separately by Mol-Graphormer are superposed based on atom mapping to construct a CGR. The feature vector of each node or edge in CGR is formed by concatenating the corresponding Mol-Graphormer embeddings in the reactant and product graphs. Missing edges resulting from bond formation or breaking are embedded as zero vectors (dashed hexagons) for concatenation. The CGR is further encoded by CGR-Graphormer. **d** Curation of the reaction-enzyme dataset. **e** VenusRXN consists of three encoders: reaction encoder, protein encoder and joint encoder. The protein encoder and joint encoder are architectural variants of the same PLM and share parameters, where the former encodes proteins only and the latter integrates protein representation with the reaction encoder output through cross-attention. The three encoders are jointly trained through cross-modal alignment of reactions and enzymes, and discriminative classification of reaction-enzyme pairs. **f** VenusRXN supports three applications: enzyme retrieval queried by reactions, enzyme retrieval queried by template enzymes, and fine-tuning on catalytic performance labels for task-specific enzyme recommendation.

To effectively learn the cross-modal interactions between reactions and enzymes, VenusRXN integrates the reaction encoder with a PLM, and jointly trains them on a large dataset of reaction-enzyme pairs through multimodal learning. The reaction-enzyme dataset is curated from BRENDA [28] and Rhea [29] databases, comprising 265,920 reaction-enzyme pairs (Fig. 1d and Methods). In VenusRXN, the PLM is modified to work in two modes: as a reaction-protein joint encoder and as a unimodal protein encoder. The joint encoder equips cross attention in its Transformer blocks to match and fuse the protein embeddings and reaction encoder output. The protein encoder shares parameters with the joint encoder, but its cross-attention layers are deactivated and it only requires protein input. During multimodal training, the embedding spaces of the reaction encoder and protein encoder are aligned with contrastive learning [25] and soft-label alignment technique [26], in which we implement a constrained batch sampling strategy to handle the many-to-many relationship between reactions and enzymes (Fig. 1e left and Methods). Meanwhile, the joint encoder is trained to classify the positive and negative reaction-enzyme pairs, where hard negatives are sampled in-batch according to contrastive similarity (Fig. 1e right and Methods). With this multi-task learning approach, VenusRXN simultaneously learns global and local matching patterns between reactions and enzymes.

VenusRXN adopts sequence-based PLM and does not rely on protein structures, so it is suitable for searching large protein sequence databases. For enzyme retrieval, the protein encoder is first used to extract embeddings for all protein sequences in the target database. These embeddings are cached to a vector database to enable rapid dense retrieval (Fig. 1f left). If the query is a reaction, the reaction encoder is applied to project it into the embedding space; or if it is a template enzyme, the protein encoder is applied. Since the embedding spaces of the reaction and protein encoders are well aligned, potential enzymes that can catalyze the query reaction or with similar catalytic functions to the template enzyme can be efficiently retrieved by computing embedding similarities between the query and the candidate proteins. Besides, since the joint encoder learns fused representations of reactions and enzymes through reaction-enzyme matching, it can be efficiently transferred to supervised learning tasks with reaction-enzyme pairs as input. Consequently, when enzyme catalytic performances for the target reactions are measured, e.g., *k*_cat_ or activity values, the joint encoder can be fine-tuned on the labeled reaction-enzyme pairs to recommend new enzymes with enhanced performances (Fig. 1f right).

### VenusRXN achieves robust reaction encoding

One approach to reaction-conditioned enzyme discovery is to predict the EC numbers of the reactions and then retrieve the enzymes associated with the predicted EC numbers [23,30]. Although VenusRXN operates independently of enzyme annotations, the EC number prediction task serves as a suitable benchmark for evaluating its reaction encoder’s ability to capture biochemical semantics in enzymatic reactions. We therefore employ this task to assess how effectively the reaction encoder can support downstream enzyme discovery workflows.

We train and test VenusRXN’s reaction encoder on the reaction EC number prediction dataset of CARE [24]. Notably, the reaction encoder evaluated here has not yet undergone multimodal learning, which means it relies solely on reaction data and has not been exposed to enzyme sequences, ensuring no data leakage from our reaction-enzyme dataset. For simplicity, we call the reaction encoder VenusRXN-RE. The CARE dataset includes three difficulty levels of train-test splits, where more challenging splits correspond to lower similarity between training and test reactions. The reaction similarity for splitting is determined by the amount of overlap in EC numbers. To overcome the data scarcity and class imbalance in EC number prediction, VenusRXN-RE is trained via supervised contrastive learning [31], which pulls reactions with the same EC number closer in the embedding space while pushing apart those with different EC numbers. During testing, a nearest-neighbor prediction approach is employed: the training reaction with the highest embedding similarity to the test reaction is retrieved, and the former’s EC number is assigned as the prediction.

The baselines include DRFP [32], CLIPZyme [21], and CREEP [23]. DRFP is an efficient reaction fingerprint calculated by set differences between product and reactant fingerprints. When using DRFP for EC number prediction, a nearest-neighbor approach based on DRFP similarity is adopted [23]. CLIPZyme and CREEP are multimodal models that leverage both reactions and enzymes for contrastive learning. Compared with VenusRXN-RE and DRFP, these two baselines adopt complicated multimodal retrieval for EC number prediction [23]. Additionally, we conduct an ablation study on VenusRXN-RE by evaluating its two variants: VenusRXN-RE (w/o CGR) excludes the CGR representation and instead concatenates the CLS embeddings of the reactants and products given by Mol-Graphormer to form the reaction embedding; VenusRXN-RE (w/o CGR & pretraining) excludes CGR, and its Mol-Graphormer is randomly initialized instead of pretrained.

Supplementary Fig. 1 shows the prediction accuracy of each model for different EC number levels and difficulties. The results for DRFP, CLIPZyme, and CREEP are obtained from CARE. As demonstrated, VenusRXN-RE obviously outperforms these baselines in almost all cases. When predicting level 4 EC numbers, it achieves a superior accuracy of 71.5%. Compared to CLIPZyme and CREEP, VenusRXN-RE is trained on unimodal data of reactions (without enzymes) and has much fewer parameters. Its superior performance thus indicates its capability to extract more discriminative and informative reaction features. When CGR is removed, VenusRXN-RE exhibits performance degradation on medium and hard difficulties. This suggests that CGR helps the model better learn reaction rules and generalize to reactions that differ greatly from the training data. Moreover, VenusRXN-RE (w/o CGR & pretraining) shows much worse performance than the original model, which confirms that the proposed pretraining strategy enables Mol-Graphormer to provide high-quality, reaction-aware molecular graph representations. In conclusion, VenusRXN-RE is a reliable reaction encoder for enzyme discovery.

### Enzyme retrieval queried by reactions

Retrieving enzymes via reaction EC number prediction confronts a major challenge: it cannot directly retrieve enzymes lacking EC annotations. In this section, we illustrate the capability of VenusRXN on straightforward, annotation-free enzyme retrieval from reaction queries. To evaluate VenusRXN’s retrieval performance on new reactions not encountered during training, we split our reaction-enzyme dataset into training and test set by substrates such that all test reactions are excluded from training data. Specifically, for each reaction in the dataset, a representative substrate is identified by first excluding common cofactors and then choosing the reactant with the largest number of atoms. Next, we randomly select 800 substrates from the representative substrate set and take all reaction-enzyme pairs corresponding to these substrates as test set, with the remaining pairs as training set. The resultant test set comprises 19,195 reaction-enzyme pairs spanning 1649 reactions and 17,096 enzymes.

During testing, with each test reaction as query, we screen all 178,035 enzymes in the dataset to identify the positive enzymes associated with it. Model performance is evaluated using four metrics: BEDROC [33], enrichment factor [34], top-*k* recall rate, and top-*k* hit rate. Top-*k* hit rate is the proportion of test queries for which at least one positive enzyme appears within the top-*k* retrieval results. To further assess generalization capability, we split the test set into subsets based on the max similarity between each test reaction and the training reactions, where the reaction similarity is measured as cosine similarity between DRFPs (the train-test reaction similarity distribution is shown in Supplementary Fig. 2). The evaluation metrics calculated on different subsets are reported.

**Fig. 2.**
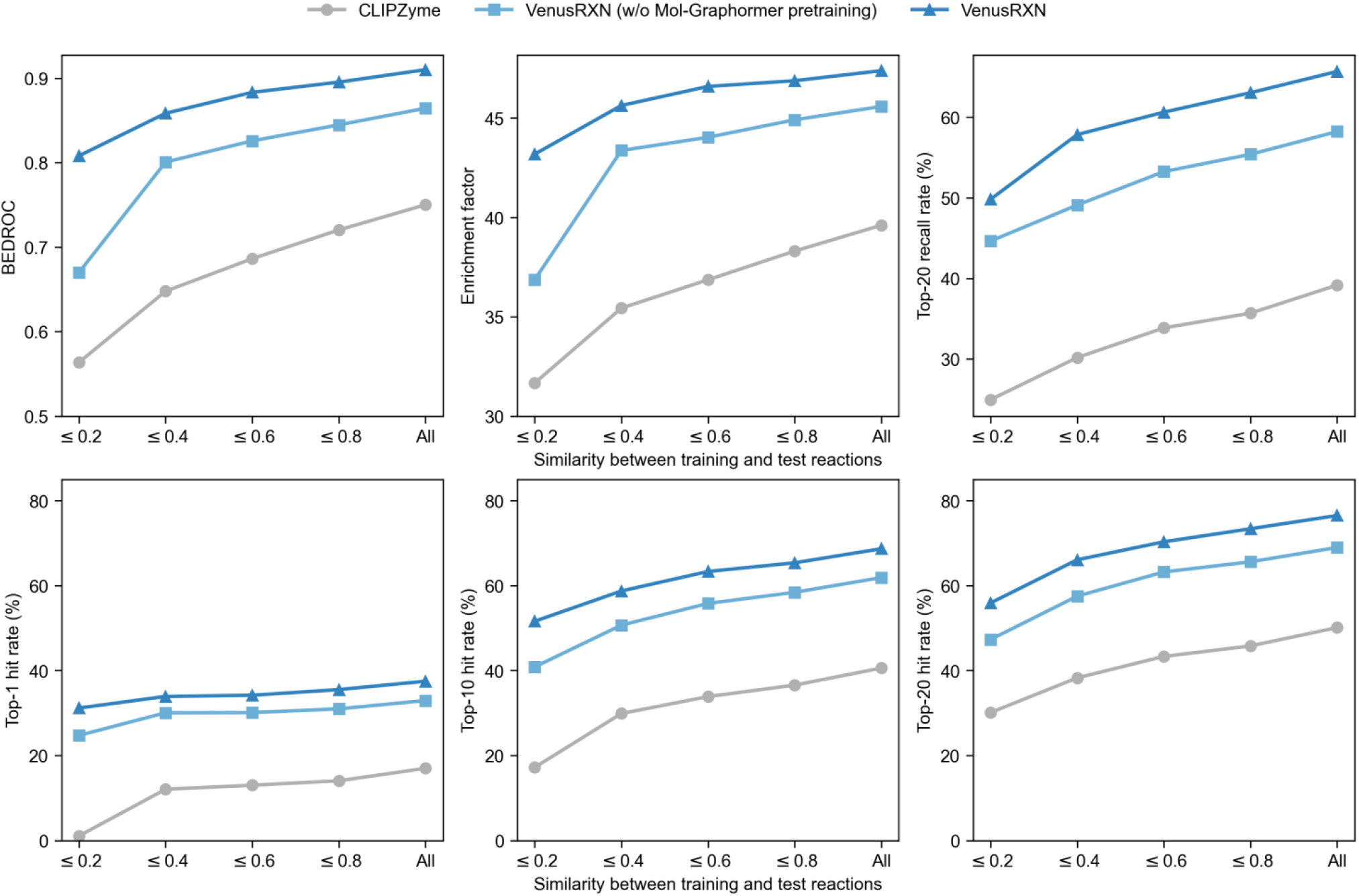
Enzyme retrieval performance when queried by reactions. The parameter *α* of BEDROC is set to 85, and enrichment factor is computed at the 2% level. We split the test set into subsets based on the max similarity between each test reaction and the training reactions, and report the evaluation metrics on different subsets. The reaction similarity for splitting is measured via cosine similarity between DRFPs.

The baseline model is CLIPZyme [21], which aligns molecular graph representations of reactions with tertiary structure representations of enzymes through contrastive learning. For fair comparison, CLIPZyme is retrained on our training set and adopts the same testing protocol as VenusRXN. Since CLIPZyme requires protein structures as input, we first attempt to download structures for all enzymes in our dataset from AlphaFold Database [35], and for enzymes not available in the database, we then predict their structures using AlphaFold2 [36]. Additionally, we evaluate VenusRXN with non-pre-trained Mol-Graphormer for ablation study, denoted as VenusRXN (w/o Mol-Graphormer pretraining).

Fig. 2 shows the retrieval performance of each model. As seen, both VenusRXN and its variant outperform CLIPZyme by a large gap across all metrics, despite not utilizing protein structures. Specifically, VenusRXN achieves a 76.5% top-20 hit rate on the full test set, surpassing CLIPZyme by 26%. Even when train-test reaction similarity drops to 0.2 or below, VenusRXN still performs well, with its top-20 recall and hit rates (49.8% and 55.9%, respectively) being nearly double those of CLIPZyme. These results demonstrate that the multimodal learning strategy of VenusRXN enables effective linking between enzymes and their catalyzed reactions as well as good generalization to reactions dissimilar from the training set. Unlike CLIPZyme which uses contrastive learning alone, VenusRXN simultaneously learns fine-grained alignment and fusion of the reaction and protein representations, enabling better capture of reaction-enzyme matching patterns and more accurate cross-modal retrieval. Besides, VenusRXN (w/o Mol-Graphormer pretraining) delivers inferior performance compared to the original model. This is consistent with the results in Supplementary Fig. 1, reaffirming the critical role of Mol-Graphormer pretraining for precise enzyme discovery.

### Enzyme retrieval queried by templates

In scenarios where known enzyme catalysts exist for a target reaction, a typical approach to discover novel enzymes is to use existing ones as templates and search for their functional analogs. To evaluate the retrieval performance of VenusRXN when queried by template enzymes, we first filter the above test set to include only reactions associated with at least two positive enzymes, leaving 872 test reactions. For each of these reactions, we then randomly select one of its positive enzymes to serve as the template. In contrast to the reaction-query experiment, here for each test reaction, the query is replaced by the selected template and the template is excluded from the retrieval database. To determine whether VenusRXN can identify functional similarity beyond sequence similarity, we split the filtered test set into subsets based on the max sequence identity between each template and its associated remaining positive enzymes (the template-positive sequence identity distribution is shown in Supplementary Fig. 3). The evaluation metrics calculated on different subsets are reported.

**Fig. 3.**
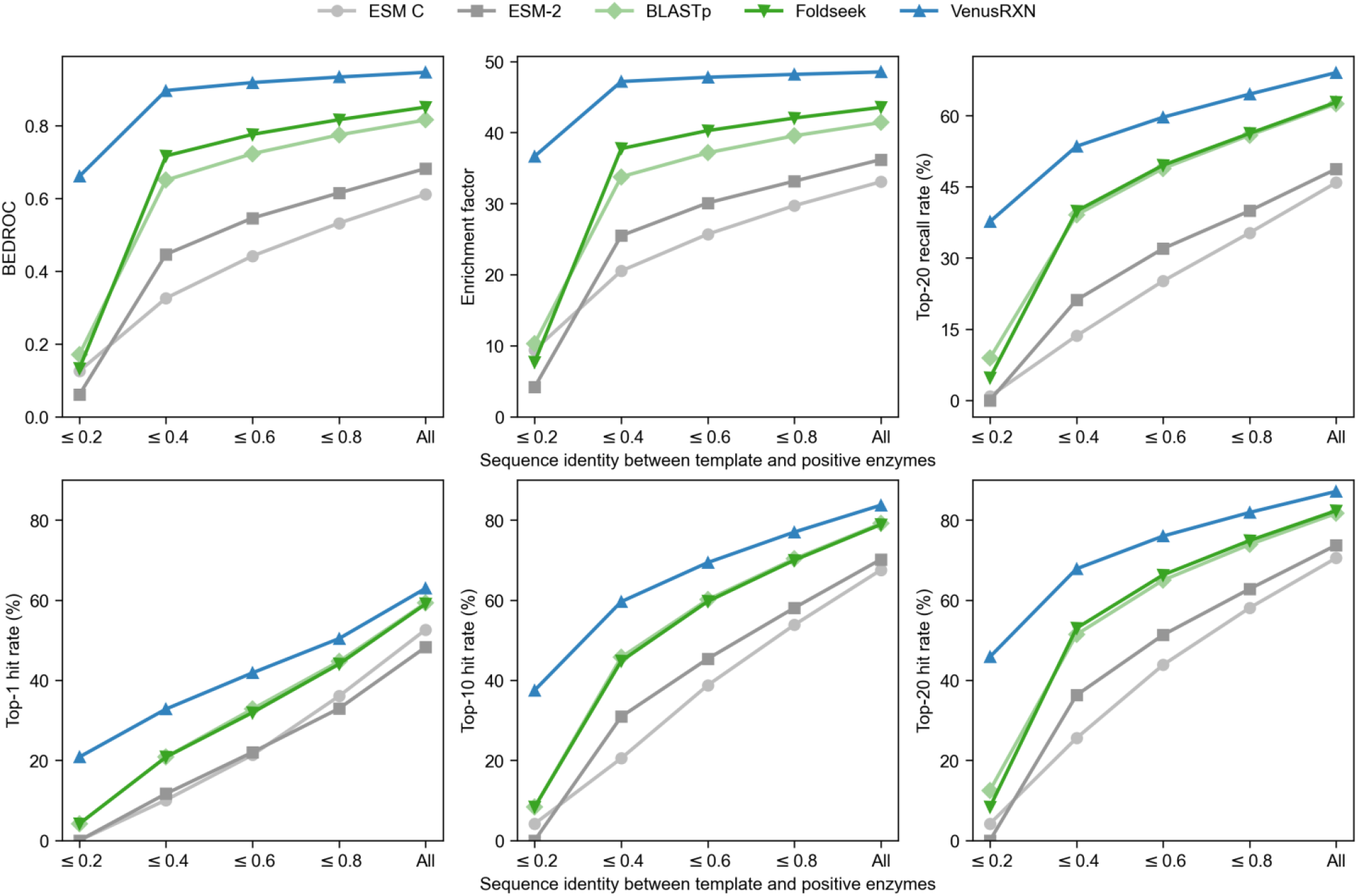
Enzyme retrieval performance when queried by template enzymes. The parameter *α* of BEDROC is set to 85, and enrichment factor is computed at the 2% level. For each test reaction, we randomly select one of its positive enzymes to serve as the template and exclude it from the retrieval database. We split the test set into subsets based on the max sequence identity between each template and its associated remaining positive enzymes, and report the evaluation metrics on different subsets.

The baseline methods include dense retrieval using raw protein embeddings from ESM-2 and ESM C, sequence similarity search via BLASTp [6], and structure similarity search via Foldseek [7]. For ESM-2 and ESM C, protein embeddings are obtained by average-pooling their top-layer embeddings. The protein structures used by Foldseek are identical to those for CLIPZyme.

The retrieval performance of each model is shown in Fig. 3. While taking only protein sequences, VenusRXN emerges as the top-performing model in all cases, surpassing both alignment- and PLM-based baselines. In particular, its advantage over the baselines is most apparent when the positive enzymes have low sequence identity (≤ 0.2) to the template, where it attains a top-20 hit rate of 45.8% versus Foldseek’s 8.3%. Its superior performance originates from its multimodal training paradigm: compared to the baselines that only leverage protein information, its protein encoder is guided and constrained by reaction context, and thus its protein embeddings can better capture catalytic functions, enabling identification of functionally similar enzymes even when their sequence or structure similarity to the template is low. On the other hand, by explicitly modeling intra-modal semantic relationships within reactions and enzymes through soft-label alignment, VenusRXN ensures robust performance in both cross-modal and intra-modal retrieval. Compared to the reaction-query results in Fig. 2, VenusRXN achieves much higher retrieval performance when taking enzyme queries, e.g., the top-20 hit rate increases from 76.5% to 87.2% on full test set. This can be primarily attributed to the fact that unimodal (enzyme-to-enzyme) retrieval is inherently less challenging than cross-modal (reaction-to-enzyme) retrieval.

### Validating VenusRXN for enzyme discovery in natural product biosynthesis

*De novo* biosynthesis of high-value metabolites represents a strategic priority for sustainable biomanufacturing. Homology-based pathway elucidation tools such as antiSMASH [37] are intrinsically limited for novel reactions absent from annotated databases. In contrast, VenusRXN enables annotation-agnostic pathway reconstruction through reaction-to-enzyme retrieval. Here, we select four recently elucidated metabolic pathways that postdate VenusRXN’s training data (after 2025.2) to evaluate its performance in reaction-to-enzyme retrieval. They include the biosynthetic pathways of Taxol [38], salicylic acid (SA) [39], sec-pentyl-TeA (S-TeA) [40], and sulfenicin [41], which present in plant, fungal, and bacterial metabolic networks, respectively. We keep only catalytic steps confirmed through *in vitro* functional validation or heterologous pathway reconstruction, resulting in 10 reactions for case study (Fig. 4). For each pathway, genome-wide protein sequences of its established host organism are obtained from NCBI [27] and JGI [42]. Then, with each of the 10 reactions as query, VenusRXN screens and ranks all protein sequences in its corresponding genome. We apply the VenusRXN model trained on our reaction-enzyme dataset directly for evaluation without any fine-tuning.

**Fig. 4.**
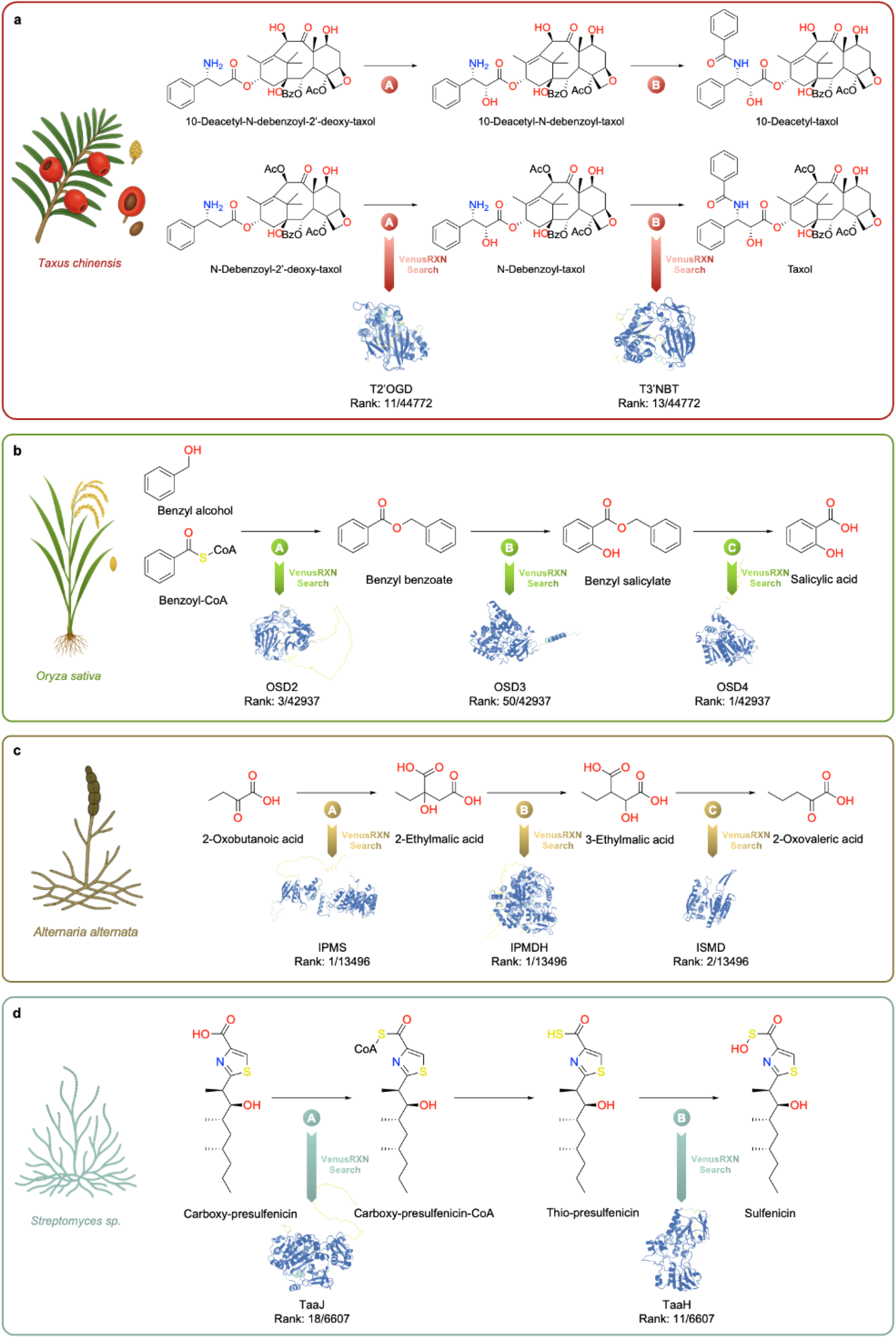
VenusRXN identifies reported enzymes from genomes in natural product biosynthesis. **a** Biosynthetic pathway of Taxol in *Taxus chinensis*: N-debenzoyl-2’-deoxytaxol is converted to Taxol via C2’α hydroxylation (by T2’OGD) and 3’-N-benzoylation (by T3’NBT). **b** Biosynthetic pathway of salicylic acid (SA) in *Oryza sativa*: Benzyl-CoA is converted to salicylic acid through benzoylation (by OSD2), hydroxylation (by OSD3), and hydrolysis (by OSD4). **c** Biosynthetic pathway of secpentyl-TeA (S-TeA) in *Alternaria alternata*: 2-Oxobutanoic acid is converted to 2-Oxovaleric acid via condensation (by IPMS), isomerization (by IPMDH), and decarboxylation (by ISMD). **d** Biosynthetic pathway of sulfenicin in *Streptomyces* sp. CNT360: Carboxy-presulfenicin is converted to sulfenicin via thioesterification (by TaaJ) and S-hydroxylation (by TaaH). The displayed enzyme structures are predicted by AlphaFold2.

Taxol is an effective natural broad-spectrum anticancer drug. Despite its clinical importance, the key enzymes responsible for the final two steps of Taxol synthesis had remained elusive for a long time. For these two challenging steps, VenusRXN ranks the target enzymes T2’OGD (for C2’α hydroxylation) and T3’NBT (for 3’-N-benzoylation) 11th and 13th respectively, among 44,772 proteins encoded by the *Taxus chinensis* genome (Fig. 4a). Even without co-expression analysis based on 59 transcriptomic datasets employed in previous study [38], VenusRXN markedly narrows the candidate pool to fewer than 15 genes.

SA is a plant hormone. The phenylalanine ammonia lyase (PAL) pathway is one of the two core routes for SA synthesis in plants, with its key enzymes and complete steps recently elucidated in *Oryza sativa*. For SA synthesis pathway, VenusRXN accurately identifies O-benzoyltransferase (OSD2) and esterase (OSD4), ranking them 3rd and 1st out of 42,937 candidates, respectively (Fig. 4b).

S-TeA is synthesized by *Alternaria alternata* and holds potential as a new synthetic herbicide. In its biosynthesis, the enzymes IPMS (for condensation), IPMDH (for isomerization), and ISMD (for decarboxylation) form a unique carbon chain elongation module. VenusRXN ranks each of them within the top 2 among 13,496 candidates (Fig. 4c).

Sulfenicin is a sulfur-containing natural product identified in *Streptomyces* sp. CNT360, featuring an acylsulfenic acid functional group. For the first step of sulfenicin biosynthesis, the target enzyme TaaJ is ranked 18th out of 6607 candidates by VenusRXN. In the last step, the S-hydroxylation reaction that forms acyl sulfenic acid was previously unknown, for which the target enzyme TaaH is ranked 11th (Fig. 4d).

Overall, VenusRXN demonstrates a high top-20 hit rate of 90% on these 10 reactions, confirming its robust enzyme retrieval performance and potential for applications in natural product biosynthesis.

### Applying VenusRXN to massive-scale, template-free enzyme mining

VenusRXN employs efficient vector similarity search for enzyme retrieval, and does not rely on protein structures, making it inherently suitable for screening large-scale sequence databases. To evaluate its large-scale screening capability, we select two challenging reactions absent from its training data— one of which involves a non-natural substrate and represents a novel reaction. With each of these two reactions as query, we screen the NCBI NR database [27] (containing over 300 million protein sequences) via VenusRXN and pick the top 10 ranked candidates for wet-lab experimental validation.

We first target the biosynthesis of (*R*)-1-cbz-3-aminopiperidine, a key chiral intermediate for dipeptidyl peptidase IV inhibitors linagliptin and alogliptin, which are frontline drugs for type 2 diabetes [43,44]. While conventional chemical synthesis relies on transition-metal catalysts that generate hazardous waste, the biosynthetic route requires an ω-transaminase (ω-TA) capable of accepting a bulky, non-natural ketone using D-serine as an amine donor (Fig. 5a). Given that this specific transformation lacks reported natural enzymes, it poses a challenge for zero-shot enzyme retrieval. Of the 10 candidates retrieved by VenusRXN, 8 are successfully expressed (Methods and Supplementary Table 1). High-performance liquid chromatography (HPLC) analysis of these expressed sequences reveals that one candidate, TA-3 (Fig. 5c), exhibits significant catalytic activity, achieving a 45% conversion rate with excellent stereoselectivity (>99% enantiomeric excess) (Fig. 5e, Methods and Supplementary Fig. 4).

**Fig. 5.**
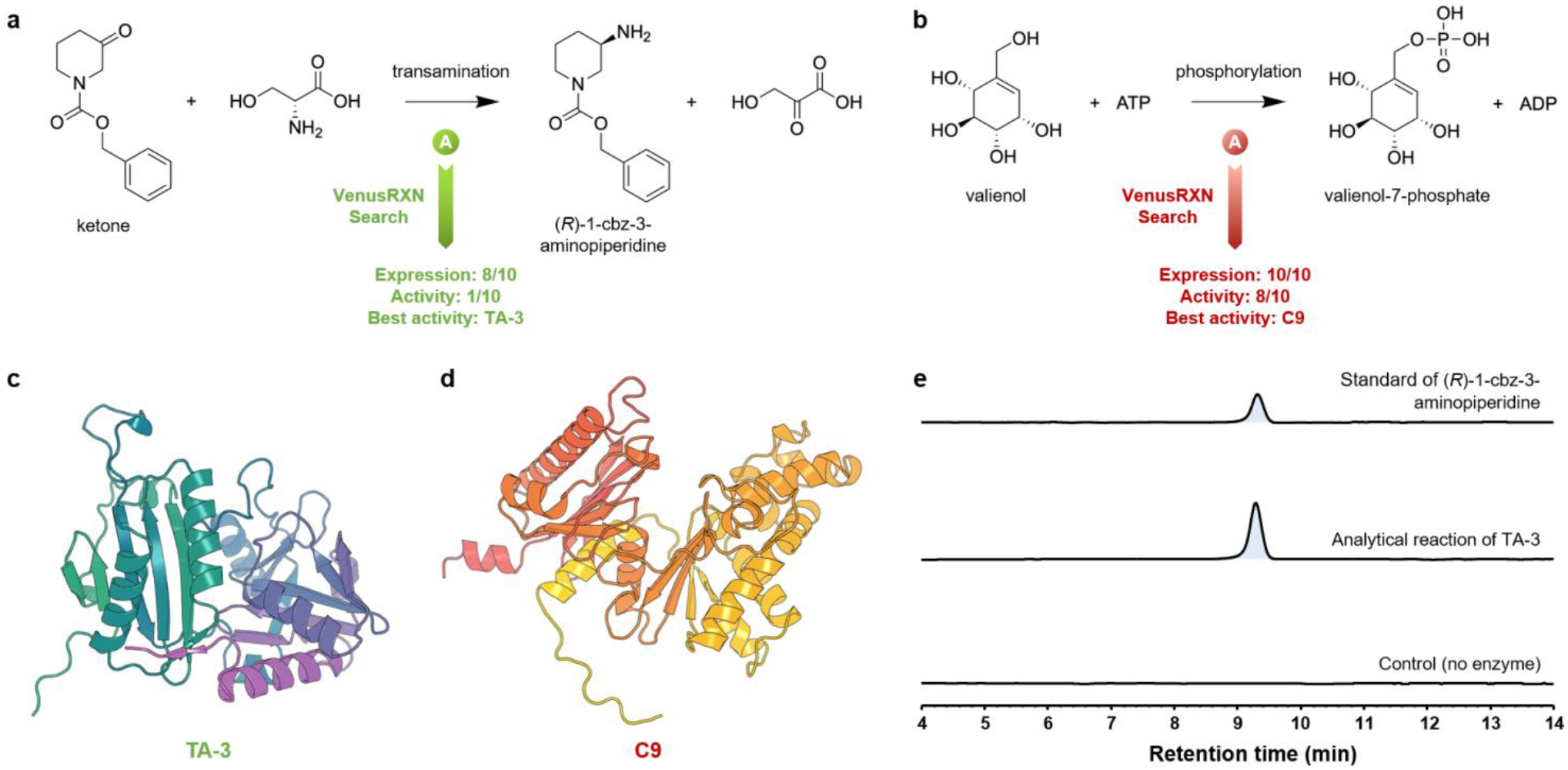
VenusRXN identifies enzymes from the NCBI NR database for transamination and phosphorylation reactions. **a** The synthesis of (*R*)-1-cbz-3-aminopiperidine, a key intermediate for diabetes drugs, from a bulky, non-natural ketone using D-serine as an amine donor. VenusRXN successfully retrieves a functional enzyme TA-3 for this reaction. **b** The phosphorylation of valienol to valienol-7-phosphate, an intermediate in acarbose biosynthesis. Among the 10 candidates retrieved by VenusRXN, C9 exhibits the best catalytic activity. **c** Predicted tertiary structure of the identified ω-transaminase TA-3. **d** Predicted tertiary structure of the identified kinase C9. **e** HPLC analysis validating the catalytic activity of TA-3.

We then validate VenusRXN on the phosphorylation of valienol to valienol-7-phosphate (Fig. 5b). Valienol-7-phosphate is one of the intermediates in acarbose biosynthesis [45], a clinically essential α-glucosidase inhibitor prescribed globally for the management of type 2 diabetes mellitus. Beyond this primary role, valienol-7-phosphate also functions as a versatile precursor for a diverse class of C_7_N-aminocyclitol natural products, including the antibiotic pyralomicin [46] and the potent trehalase inhibitor salbostatin [47]. Liquid chromatography-mass spectrometry (LC-MS) analysis of cell-free reactions reveals that 8 of the 10 retrieved candidates (Supplementary Table 2) exhibit catalytic activity, with C9 showing the highest conversion efficiency (Fig.5d, Methods and Supplementary Fig. 5).

These results demonstrate the practical capability of VenusRXN to distinguish functional enzymes within a massive search space, even for query reactions with non-natural substrates.

### Fine-tuning VenusRXN to predict *k*_cat_

While determining whether an enzyme can catalyze the target reaction is crucial, enzyme discovery further strives to identify enzymes with enhanced performance. Through fine-tuning with experimental data, VenusRXN can be efficiently adapted for predicting enzyme catalytic performance (Fig. 1f right). We illustrate this by fine-tuning VenusRXN’s joint encoder to predict *k*_cat_. The *k*_cat_ dataset of TurNup [48] is used for benchmark, which consists of 4271 reaction-enzyme-*k*_cat_ triplets, covering 2977 unique reactions and 2827 unique enzymes. The data is split into training and test sets in an 8:2 ratio, with enzymes in the test set excluded from the training set. Following TurNup, all *k*_cat_ labels are log_10_-transformed to approximate a Gaussian distribution before training and testing. The joint encoder is trained using mean squared error (MSE) loss, with its matching head from multimodal learning directly repurposed as a regression head without additional parameters. The learning rate and number of epochs are determined according to 4-fold cross-validation performance on the training set.

The baseline model is TurNup, which takes DRFPs of reactions and ESM-1b [8] embeddings of enzymes as input to train an XGBoost regressor for *k*_cat_ prediction. Following TurNup, we additionally evaluate the *k*_cat_ prediction performance of VenusRXN using either reaction or enzyme input alone. For reaction-only input, we add a regression head to its reaction encoder for fine-tuning and testing; for enzyme-only input, we similarly fine-tune and test its protein encoder. Evaluation metrics include MSE, coefficient of determination (*R*^2^), Pearson correlation coefficient, and Spearman’s rank correlation coefficient.

Fig. 6 shows the prediction performance of each model. As demonstrated, regardless of the input modality, VenusRXN significantly outperforms TurNup across all metrics, achieving Spearman correlation coefficients exceeding 0.6. Notably, using only enzymes as input, the performance of VenusRXN in terms of Spearman correlation approaches that of TurNup with joint input (0.63 versus 0.65). This demonstrates that through multimodal learning, all three encoders of VenusRXN can be efficiently transferred to the *k*_cat_ prediction task and successfully identify enzymes with high *k*_cat_ values. Furthermore, for both VenusRXN and TurNup, taking reaction-enzyme joint input yields substantially better performance than taking either reaction or enzyme input alone. This result is consistent with expectations since an enzyme’s kinetic parameters depend not only on the enzyme itself but also on the catalyzed reaction, particularly the specific substrate involved. In conclusion, the joint encoder of VenusRXN effectively integrates reaction and enzyme representations through cross-attention, making it suitable for learning to predict enzyme functional properties related to enzymatic reactions.

**Fig. 6.**
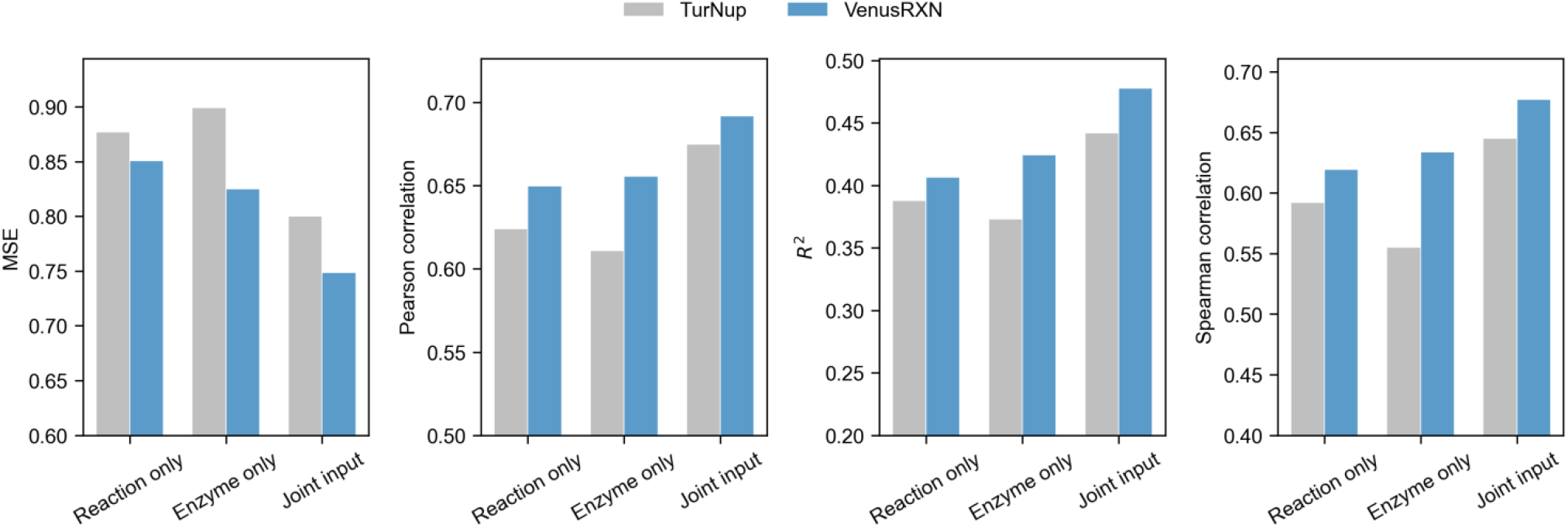
Performance on *k*_cat_ prediction. The evaluation metrics are calculated using log_10_-transformed *k*_cat_ labels. For VenusRXN, the performances of Reaction only, Enzyme only, and Joint input are obtained through fine-tuning its reaction encoder, protein encoder, and joint encoder respectively.

## Discussion

Conventional enzyme discovery methods, which primarily rely on sequence/structure homology or functional annotations, are limited in accuracy and can hardly retrieve enzymes for orphan reactions. To overcome this limitation, we propose VenusRXN, a multimodal deep learning framework for reaction-conditioned enzyme discovery. It equips a graph-based reaction encoder that utilizes self-supervised pretraining and CGR to accurately represent reactions from the atomic scale. To better learn reaction-enzyme mapping, a large and diverse dataset of reaction-enzyme pairs is curated. By jointly training the reaction encoder with a PLM on this dataset through multimodal learning, VenusRXN establishes a high-quality semantic alignment between chemical transformations and protein sequences. It covers three use cases: zero-shot reaction-to-enzyme retrieval, zero-shot enzyme-to-enzyme retrieval, and fine-tuning for task-specific enzyme recommendation. Through comprehensive benchmarks on these tasks, VenusRXN considerably outperforms prior state-of-the-art methods, and generalizes well to test reactions highly dissimilar to training reactions.

A key practical advantage of VenusRXN is its capability to efficiently screen large-scale sequence databases directly from reactions. This stems from two design features. First, it learns and predicts reaction-enzyme mapping independently of functional annotations, protein structures, and pocket information, which are typically expensive to obtain at scale. Second, it performs enzyme retrieval by encoding reactions or enzymes into fixed-length embedding vectors for alignment-free comparison. Once the vector database is built, VenusRXN allows sub-second retrieval over millions of candidates via optimized indexing, being much faster than conventional alignment- or classification-based approaches. This capability is further validated by wet-lab experiments: VenusRXN successfully identifies functional enzymes from the NCBI NR database for two challenging reactions, one of which is an unreported reaction with a non-natural substrate.

With its ability in accurately discovering enzymes for individual reactions, VenusRXN can rebuild the workflow of biosynthetic pathway elucidation. Traditional pathway elucidation approaches rely on labor-intensive methodologies: multi-omics integration (metabolomics/proteomics), gene co-expression analysis, and targeted mutagenesis, while homology-based tools (e.g. antiSMASH [37]) can hardly extrapolate to novel reactions not present in annotated databases. In comparison, using confirmed or inferred reactions as queries, VenusRXN can directly search through the genome coding sequences to suggest possible candidate genes. In our case study, VenusRXN demonstrates a high top-20 retrieval hit rate of 90% on the four newly discovered biosynthetic pathways. Therefore, with VenusRXN, future *in silico* biosynthetic pathway exploration could be more accurate and efficient, thereby largely narrows the candidate pool for wet-lab validation.

By combining its zero-shot retrieval and transfer learning capability, VenusRXN offers a scalable and unified solution for enzyme discovery. Researchers can first perform zero-shot retrieval based on either reactions or template enzymes to broadly screen candidate enzymes. If available, catalytic performance labels (e.g., *k*_cat_ and activity values) can be used to fine-tune the joint encoder of VenusRXN, such that the fine-tuned model can then re-rank the obtained candidates to prioritize the ones with enhanced performance. Future research could integrate the high-throughput screening power of VenusRXN with the structural modeling capability of geometric deep learning models to achieve more accurate enzyme discovery. It is also promising to integrate VenusRXN with state-of-the-art bio-retrosynthesis models for biosynthetic pathway design. In addition, the reaction-to-reaction and enzyme-to-reaction search capabilities supported by VenusRXN—although not explored in this study—could also be useful for characterizing novel reactions and enzymes.

## Methods

### Dataset curation

#### Reaction-enzyme dataset

To build the reaction-enzyme dataset, we extract enzymes and the reactions they catalyze from BRENDA [28] and Rhea [29] databases. The data in these two databases are curated from literature by experts, ensuring accurate reaction-enzyme annotations. The versions of the raw data downloaded from BRENDA and Rhea are 2024.1 and 2025.2.5, respectively. For the raw data of BRENDA, reactions are given as text strings where substrates and products are specified by trivial names, and enzymes are given as UniProt IDs. We follow the reaction processing pipeline of EnzymeMap [49] to resolve, standardize and correct the raw BRENDA reactions, turning them into valid SMILES [50] strings. The corresponding enzyme sequences are then downloaded from UniProt [1]. For Rhea, the reaction SMILES and enzyme sequences are already provided in the raw data. We use RXNMapper [51] to atom map all reactions. If a reaction is unbalanced, we further try the heuristics provided by EnzymeMap to correct and atom map it. VenusRXN utilizes atom mapping to identify the reaction center and molecular transformations in a reaction. After filtering out reactions that fail to be atom mapped, merging and deduplicating reaction-enzyme pairs from both databases, we finally obtain a total of 265,920 reaction-enzyme pairs, comprising 23,595 unique reactions and 178,035 unique enzymes.

#### Reaction dataset for Mol-Graphormer pretraining

The pretraining reaction data for Mol-Graphormer is curated by merging all reactions from our reaction-enzyme dataset with the reactions in USPTO [52] and CARE [23] datasets. For reactions lacking atom mappings in USPTO and CARE, the aforementioned atom mapping process is applied. After deduplication, a final dataset of 1,145,180 reactions is obtained.

### Reaction encoder

VenuxRXN’s reaction encoder consists of two Graphormer models: Mol-Graphormer and CGR-Graphormer. Mol-Graphormer is pretrained on our reaction dataset through self-supervised learning and provides basic molecular features of reactants and products. Based on the output embeddings of Mol-Graphormer, CGR-Graphormer further captures the molecular transformations in a reaction by encoding its CGR.

#### Mol-Graphormer architecture

Given a reaction SMILES, the molecular graphs of reactant(s) and product(s) are constructed using RDKit [53], where the Hs are implicit in the graphs. Supplementary Table 3 and 4 show the node (atomic) features and edge (bond) features of the molecular graphs respectively. On either side of the reaction, if multiple molecules exist, their molecular graphs are combined into a single disconnected graph, allowing Mol-Graphormer to model interactions between different reactant or product molecules through attention mechanism. The reactant graph and product graph are then fed into Mol-Graphormer separately. Following the original Graphormer [54], Mol-Graphormer extends the standard self-attention in Transformer by spatial encoding and edge encoding to learn the molecular structures. Specifically, the attention score between nodes *v*_*i*_ and *v*_*j*_ is computed as follows:

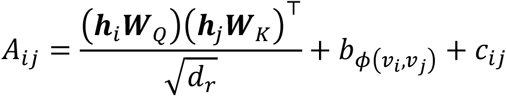

where ***h*** denotes node embedding vector, ***W*** denotes learnable weight matrix, *b* and *c* are learnable scalars describing the graph topology, and *d*_*r*_ is the hidden size of Mol-Graphormer. The index function *ϕ*(*v*_*i*_, *v*_*j*_) is defined by the shortest path length *D*_*ij*_ between nodes *v*_*i*_ and *v*_*j*_ in the molecular graph:

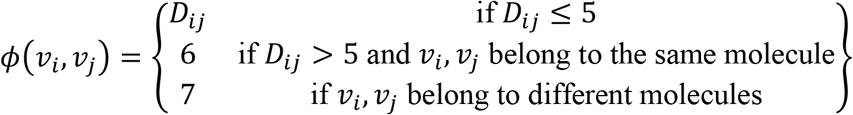

*c*_*ij*_ is computed based on the edge features along the shortest path between *v*_*i*_ and *v*_*j*_:

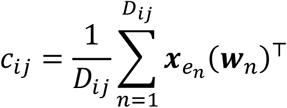

where 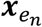 denotes the feature vector of the *n*-th edge *e*_*n*_ on the shortest path, and ***w***_*n*_ is the *n*-th weight vector. Besides, like the original Graphormer [54], a virtual node connected to all existing nodes is added to the graph and its top-layer embedding is served as a graph-level readout. Imitating BERT [55], we call them CLS node and CLS embedding. With Mol-Graphormer, the long-range node interactions within molecular graphs can be better modeled compared with traditional message-passing neural networks. The hyperparameters of Mol-Graphormer are detailed in Supplementary Table 5.

#### Mol-Graphormer pretraining

The pretraining of Mol-Graphormer is performed on both reactant and product graphs (Fig. 1b). We design three self-supervised learning tasks: MAP, RCP, and GCL.

In the MAP task, a certain proportion of the atoms in both reactant and product graphs are randomly masked out, and the model must recover the properties of these atoms given the rest of the corresponding graph. This task draws inspiration from the masked language modeling objective of BERT and aims to force the model to learn contextualized representations of graph nodes, thereby enhancing its capability to model local graph structures. Since the atomic feature values in the dataset are highly unbalanced, we combine features 1, 3, 4, 5 in Supplementary Table 3 into a composite property C1, and features 1, 2, 3, 8, 9 into C2 for prediction to increase task difficulty. Specifically, the output embedding of each masked atom is fed into a multi-layer perceptron (MLP) to predict the probability distribution over possible property values. Cross-entropy loss is then adopted for training:

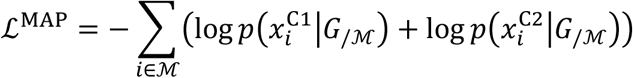

where ℳ denotes the index set of the masked atoms, *G*_/ℳ_ denotes the masked reactant or product graph, 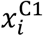 and 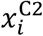 are the true property values of the *i*-th atom. The masking scheme follows BERT: 15% of the atoms in each graph are randomly selected for masking; for each selected atom, 80% of the time we replace it with a MASK token, 10% of the time we replace its input with a random feature vector, and 10% of the time we leave it unchanged.

In the RCP task, the model must first identify reaction center atoms (atoms whose properties or bonds change during the reaction) in the reactant and product graphs respectively, and then predict their properties on the other side of the reaction. This task forces the model to localize key atoms and functional groups involved in the reaction, and thus helps it infer general reaction rules. Similarly, the atom embeddings are fed into an MLP to perform classification tasks:

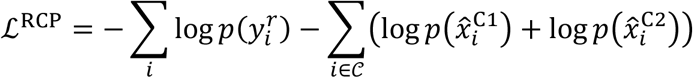

where 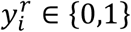 denotes the true reaction center label of the *i*-th atom, 𝒞 is the index set of the reaction center atoms, 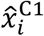 and 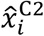 are atom *i*’s true property values on the other side of the reaction. The prediction targets 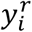 and 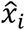 in this task are obtained by atom mapping.

In the GCL task, reactant-product graph pairs capable of forming a reaction are treated as positive pairs, while those incapable of are considered negative pairs, and the model is trained to distinguish between the positives and negatives (i.e., valid and invalid reactions) in its embedding space. This task forces the model to extract representative global features of molecular graphs, and facilitates it to learn the relations between reactants and products. Specifically, the CLS embeddings of reactant and product graphs are first projected into a shared low-dimensional embedding space with different MLPs for comparison. Afterwards, the InfoNCE [56] loss is applied for training. Assuming a training batch contains *N* reactions, and the reactant-product pair of the *i*-th reaction in the projected embedding space is (***u***_*i*_, ***v***_*i*_), the loss for this pair is defined as:

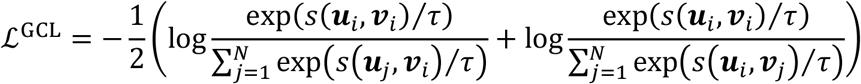

where *s* is cosine similarity function and *τ* is a learnable temperature parameter.

The final pretraining loss for Mol-Graphormer on a reaction sample is:

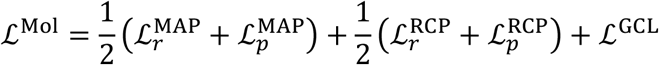

where ℒ_*r*_ and ℒ_*p*_ denote the losses computed on the reactant graph and product graph, respectively. For computational efficiency, the masked graphs constructed for the MAP task are also model input for both the RCP and GCL tasks. In this way, the reactant and product graphs only need to undergo a single forward pass per training iteration, and the masked input can be considered as a form of data augmentation for RCP and GCL. Notably, we use the same Mol-Graphormer but different output MLPs for reactants and products, and the parameters of the output MLPs are not shared across tasks. We pretrain Mol-Graphormer on 4×H100 GPUs using distributed data parallel (DDP). AdamW optimizer is applied with an initial learning rate of 0.0001, and the total batch size is 256. Gradients are clipped to a maximum norm of 5.0 to avoid gradient exploding. We randomly sample 5000 reactions from our reaction dataset as validation data and pretrain Mol-Graphormer for up to 100 epochs. After pretraining, the checkpoint with the lowest validation loss is picked.

#### Representing reactions with CGR

In cheminformatics, CGR [24] is found to be a good reaction representation for various tasks including structure-reactivity modeling [57], reaction condition prediction [58], and reaction similarity searches [59]. It is a superposition of the reactant and product graphs in a reaction, where the nodes of the two graphs are aligned by atom mapping (Fig. 1c). In this work, the feature vector of each node or edge in CGR is formed by concatenating the Mol-Graphormer embeddings of the corresponding mapped nodes or edges in the reactant and product graphs. If an edge only occurs on one side of the reaction (indicating bond formation or breaking), the edge embedding on the other side is padded with zeros before concatenation. CGR-Graphormer has similar architecture to Mol-Graphormer but it has much fewer parameters and is not pretrained. Its hyperparameters are detailed in Supplementary Table 5. In the downstream tasks, the two Graphormers are trained together.

Generally, to build the CGR of a reaction, correct atom mapping is required and the reaction is expected to be balanced. However, many reactions in the dataset are unbalanced although correction heuristics are applied. For such reactions, the CGR is constructed using only the mapped atoms in reactants and products, i.e., it is composed of subgraphs from the reactant and product graphs. In this case, although the CGR cannot cover the complete input reaction, its node features retain global reaction context captured by the attention mechanism of Mol-Graphormer. This makes the reaction encoder robust to unbalanced input reactions.

### Multimodal learning of VenusRXN

VenusRXN consists of three encoders: reaction encoder, protein encoder and joint encoder. The protein encoder and joint encoder are architecturally variant implementations of the same PLM and share parameters, where the former encodes proteins only and the latter integrates protein representation with the reaction encoder output. The three encoders are jointly trained through cross-modal alignment of reactions and enzymes, and discriminative classification of reaction-enzyme pairs (Fig. 1e).

#### Cross-modal feature alignment

We employ two complementary strategies, namely contrastive learning and soft-label alignment, to align the embedding spaces of reaction and protein encoders. This enables direct comparison between these two types of embeddings, facilitating both high-speed dense retrieval and cross-modal feature fusion. During training, the CLS embeddings produced by the reaction and protein encoders are projected into a shared low-dimensional embedding space with different linear layers (projection heads; Fig. 1e left).

The contrastive learning approach follows CLIP [25], which pulls matched reaction-enzyme pairs closer and pushes unmatched pairs apart in the projected embedding space. Formally, given a training batch containing *N* reaction-enzyme pairs, let (***r***_*i*_, ***e***_*i*_) denote the projected embeddings of the *i*-th pair. We compute the normalized reaction-to-enzyme and enzyme-to-reaction similarity scores as:

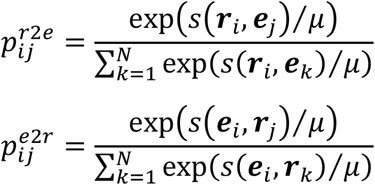

where *μ* is a learnable temperature parameter. Then, the loss for the *i*-th pair is:

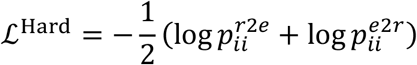

Notably, CLIP loss requires the pairs {(***r***_*i*_, ***e***_*j*_)[1 ≤ *i* ≤ *N*, 1 ≤ *j* ≤ *N, i* ≠ *j*} to be negative. However, since an enzymatic reaction can typically be catalyzed by multiple enzymes and a single enzyme can also catalyze various reactions, this requirement is difficult to meet when batches are sampled without constraints. To address this, we implement a constrained sampling strategy: for each batch, the reaction-enzyme pairs are sampled from the training set one at a time; for each sampled pair, we exclude all enzymes associated with the selected reaction and all reactions associated with the selected enzyme from the training set for subsequent sampling. The training set is not reset for each batch but only when exhausted. If it is exhausted before a batch reaches size *N*, we restart the sampling of this batch with the reset training set.

Contrastive learning can be considered as a hard-label alignment approach because it simply treats unmatched reaction-enzyme pairs as negatives, i.e., it necessitates the complete mutual exclusivity between any two unpaired samples. As a result, potential weak correlations among unmatched pairs are ignored. To mitigate this limitation, we further adopt the soft-label alignment approach [26]. It defines softened targets for cross-modal alignment based on intra-modal semantic relevance, enabling finer alignment between the two modalities. Specifically, we first compute normalized intra-modal similarity scores for both reactions and enzymes within the batch:

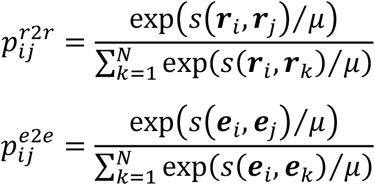

When reaction and enzyme embedding spaces are well-aligned, the intra-modal and cross-modal similarity distributions should exhibit consistency. Therefore, Kullback-Leibler (KL) divergence is used to compute the soft alignment loss:

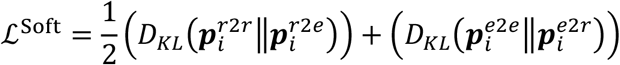

The final cross-modal alignment loss is:

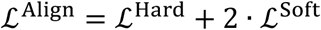

#### Cross-modal feature fusion

In the above process, reactions and enzymes are encoded as global embedding vectors ***r*** and ***e***. Despite of enabling rapid dense retrieval, this may prevent the model from learning local patterns in the input. In fact, except for CLS embeddings, the reaction encoder can provide atom-level embeddings and the PLM can provide residue-level embeddings. To better match and fuse these informative embeddings, the joint encoder equips cross-attention layers between the selfattention and feed forward layers in its Transformer blocks (Fig. 1e right). Formally, given atom embeddings ***H***_*r*_ from the reaction encoder and residue embeddings ***H***_*e*_ from the joint encoder, the cross attention is computed as:

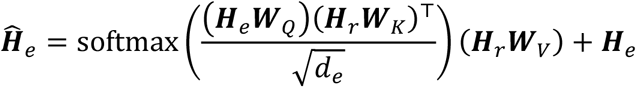

where ***W*** denotes learnable weight matrix, *d*_*e*_ is the hidden size of the PLM.

The joint encoder generates fused multimodal representation for a given reaction-protein pair. On the top of it, its CLS embedding is fed into an MLP (matching head) to compute the matching score *z* of the pair. During training, the joint encoder performs binary classification to determine whether a reaction-enzyme pair matches, with the loss function defined as:

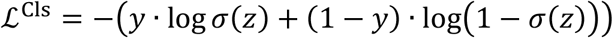

where *y* ∈ {0,1} denotes the true label of the given pair, *σ* is the sigmoid activation function. We sample hard negatives dynamically from each training batch for binary classification. For each reaction in the batch, an unmatched enzyme is sampled from the same batch according to the contrastive similarity distribution 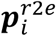 to form a negative pair. Analogously, for each enzyme, an unmatched reaction is sampled from distribution 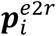 to construct a negative pair. With this strategy, the unmatched reaction-enzyme pairs that are proximate in the current embedding space are preferred to be training samples, thereby enhancing the model’s discriminative capability.

The final training loss for VenusRXN is:

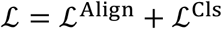

The joint encoder and protein encoder are initialized from ESM-C (600M version), with cross-attention layers inserted in its last 18 Transformer blocks. Multimodal training is conducted on 8×H200 GPUs using DDP and BFloat16 precision. We employ AdamW optimizer with an initial learning rate of 0.0001. Gradients are clipped to a maximum norm of 5.0. The total batch size is 208, and the training lasts for 400K steps.

### Validating the catalytic activity of the ω-TA

The candidate ω-TAs are expressed in *Escherichia coli* BL21(DE3) and purified via Ni-affinity chromatography. The transamination of ketone is evaluated under the following conditions: 100 µL volume with 10 mM ketone, 100 mM D-serine, 0.2 mM pyridoxal 5’-phosphate and 1 mg/mL ω-TAs in 100 mM Tris-HCl Buffer (pH 7.5), 30°C for 1 h. The reactions are quenched by adding 200 μL methanol and analyzed by precolumn derivative HPLC to determine the concentration of products. HPLC analysis is performed using an Agilent 1290 Infinity II LC System equipped with a Kromasil^®^ 100-3.5-C18 column (4.6 × 150 mm, 5 μm). Analytic program: flow rate of 1 mL/min, room temperature, H_2_O (containing 0.1% formic acid): acetonitrile = 40:60, detection wavelength of 332 nm. Chiral HPLC analysis is performed with a CHIRALPAK^®^ IG column (4.6 × 150 mm, 5 μm). Analytic program: flow rate of 0.5 mL/min, column temperature of 25°C, detection wavelength of 332 nm, mobile phase used 20–80% acetonitrile (0–15 min), 80% acetonitrile (15–25 min) [60].

### Validating the catalytic activity of the valienol kinase

#### Plasmid construction and expression

Codon-optimized genes for the retrieved candidate enzymes are cloned into the pET-30a (+) vector between the 5’ EcoRI and 3’ HindIII restriction sites, and transformed into *Escherichia coli* BL21(DE3). Cultures are grown in LB medium supplemented with 50 µg/mL kanamycin at 37°C until an OD600 nm of 0.6–0.8 is reached. Protein expression is induced with 0.4 mM IPTG (isopropyl β-D-1-thiogalactopyranosid) at 16°C for 12–16 h. Cells are harvested by centrifugation, resuspended in buffer A (20 mM Tris-HCl, pH 8.0, 300 mM NaCl), and lysed by sonication.

#### Cell-free enzymatic assays

A 30 µL reaction mixture is assembled with 25 mM Tris-HCl (pH 7.0), 10 mM MgCl_2_, 20 mM ATP, 2.5 mM valienol, and 4.33 µM enzyme (calculated by converting the crude enzyme solution concentration and the gray value ratio of the target protein). The mixture is incubated at 30°C for 6 h. A blank control is established by replacing the enzyme with an equal volume of ultrapure water. The reaction is stopped by adding two volumes of methanol, followed by vigorous vortex to denature the proteins. The mixture is centrifuged at 13,800×g for 20 min, and the supernatant is subjected to LC-MS analysis.

#### LC-MS analysis

The enzymatic reaction mixtures and related compounds are analyzed by HPLC-time-of-flight/mass spectrometry (HPLC-TOF/MS) (Agilent 1290-6230 Q-TOF) with Agilent ZORBAX SB-Aq (4.6 × 150 mm, particle size 5 μm) at a flow rate of 0.4 mL/min using an elution buffer composed of (A) Milli-Q water (containing 10 mM ammonium formate) and (B) methanol with isocratic elution at 0.5% B. Negative mode electrospray ionization is used for related compounds detection.

## Data availability

The reaction and enzyme data for training VenusRXN is obtained from USTPO [52], BRENDA [28], and Rhea [29]. The dataset of reaction EC number prediction is from CARE [24]. The dataset of *k*_cat_ prediction is from TurNup [48]. Enzyme structures in this study are downloaded from AlphaFold Database [35] or predicted by AlphaFold2 [36]. Genome-wide protein sequences of *Taxus chinensis, Oryza sativa, Alternaria alternata*, and *Streptomyces* sp. CNT360 are obtained from NCBI [27] and JGI [42].

## Code availability

The source code of VenusRXN is available at https://github.com/zy-zhou/VenusRXN.

## Acknowledgements

This work was supported by Shanghai Municipal Science and Technology Major Project, the National Key Research and Development program of China (2024YFA0917603), the Computational Biology Key Program of Shanghai Science and Technology Commission (23JS1400600), Shanghai Municipal Education Commission (2024AIZD015), Shanghai Jiao Tong University Scientific and Technological Innovation Funds (21X010200843), and Science and Technology Innovation Key R&D Program of Chongqing (CSTB2022TIADSTX0017, CSTB2024TIAD-STX0032), National Natural Science Foundation of China (62506226).

## Author contributions

P.T., and L.H. conceptualized and supervised this research project. Z.Z., Y.H. and P.T. developed the methodology. Z.Z. and Y.H. implemented the method and performed the benchmark. Y.Z. and X.F. conducted the wet-lab experiments. Z.Z, B.Z., X.C., Q.X, J.H., R.H., S.L., L.B. and P.T. wrote the manuscript. All authors reviewed and accepted the manuscript.

